# Efficacy of longevity interventions in *C. elegans* is determined by early life activity of RNA splicing factors

**DOI:** 10.1101/2021.11.01.466772

**Authors:** Sneha Dutta, Caroline Heintz, Maria C. Perez-Matos, Ayse Sena Mutlu, Mary E. Piper, Meeta Mistry, Arpit Sharma, Hannah J. Smith, Porsha Howell, Rohan Sehgal, Anne Lanjuin, Meng C. Wang, William B. Mair

## Abstract

Geroscience aims to target the aging process to extend healthspan. However, even isogenic individuals show heterogeneity in natural aging rate and responsiveness to pro-longevity interventions, limiting translational potential. Using *in vivo* mini gene reporters in isogenic *C. elegans*, we show that alternative splicing of mRNAs related to lipid metabolism in young animals is coupled to subsequent life expectancy. Further, activity of RNA splicing factors REPO-1 and SFA-1 early in life modulates effectiveness of specific longevity interventions via POD-2/ACC1 and regulation of lipid utilization. In addition, early inhibition of REPO-1 renders animals refractory to late onset suppression of the TORC1 pathway. Together these data suggest that activity of RNA splicing factors and the metabolic landscape early in life can modulate responsiveness to longevity interventions and may explain variance in efficacy between individuals.

**One Sentence Summary:** Efficacy of pro-longevity interventions in *C. elegans* is determined by the activity of splicing factors and the lipid metabolic landscape early in the life of the individual.

---

Aging is the key risk factor for most non-communicable chronic diseases. In the last twenty years the field of geroscience has discovered genetic, metabolic and pharmacological pro-longevity interventions that delay aging in multiple species and might be used to promote healthspan in the elderly. These interventions include dietary restriction (DR), intermittent fasting (IF), and genetic or pharmacological modulation of key metabolic and stress response pathways (*1*). However, the effectiveness of these interventions is highly variable between individuals, sexes and genotypes (*2*). Given the greater heterogeneity in humans compared to laboratory model organisms, such variance currently limits effective translation of geroscience discoveries to usable therapeutics for the elderly. Understanding the biological mechanisms that underpin heterogeneity of treatment response is therefore needed if we are to leverage the discovery of anti-aging regimens to alleviate diseases of aging in people.

## RESULTS

Deregulation of pre-mRNA splicing has been implicated in multiple age-related chronic diseases (*3*), and changes in expression of key splicing regulators and splicing of their targets are associated with longevity in mice and humans (*4, 5*). Moreover, specific RNA splicing factors have been shown to be causally linked to the effects of pro-longevity interventions in *C. elegans* (*6*–*8*). We therefore hypothesized that changes in alternative splicing of specific pre-mRNAs might be coupled to both the rate of biological aging and response to longevity interventions. To test this, we first sought to define early RNA processing events that correlate with subsequent life expectancy. Fluorescent mini gene reporters of a single exon 5 skipping event in *ret-1* (*9*) in *C. elegans* can be used to sort isogenic animals at day 6 of adulthood into subpopulations with short or long life-expectancies, identifying animals that are naturally aging poorly or well (*6*). Animals with prevalent *ret-1* exon 5 inclusion (GFP) at day 6 show lifespan extension compared to isogenic animals with exon 5 skipping (mCherry), defined therefore as having long life expectancy (LL) versus short life expectancy (SL) respectively (Fig 1A-C). We isolated LL or SL subpopulations at day 6 day in sextuplicate (Fig 1C, S1A) and performed 75-bp paired-end RNA-Seq followed by analysis of gene expression and differential isoform usage. This allowed us to define genes in which either expression level or isoform usage differed between LL v SL animals.

**Figure 1:**
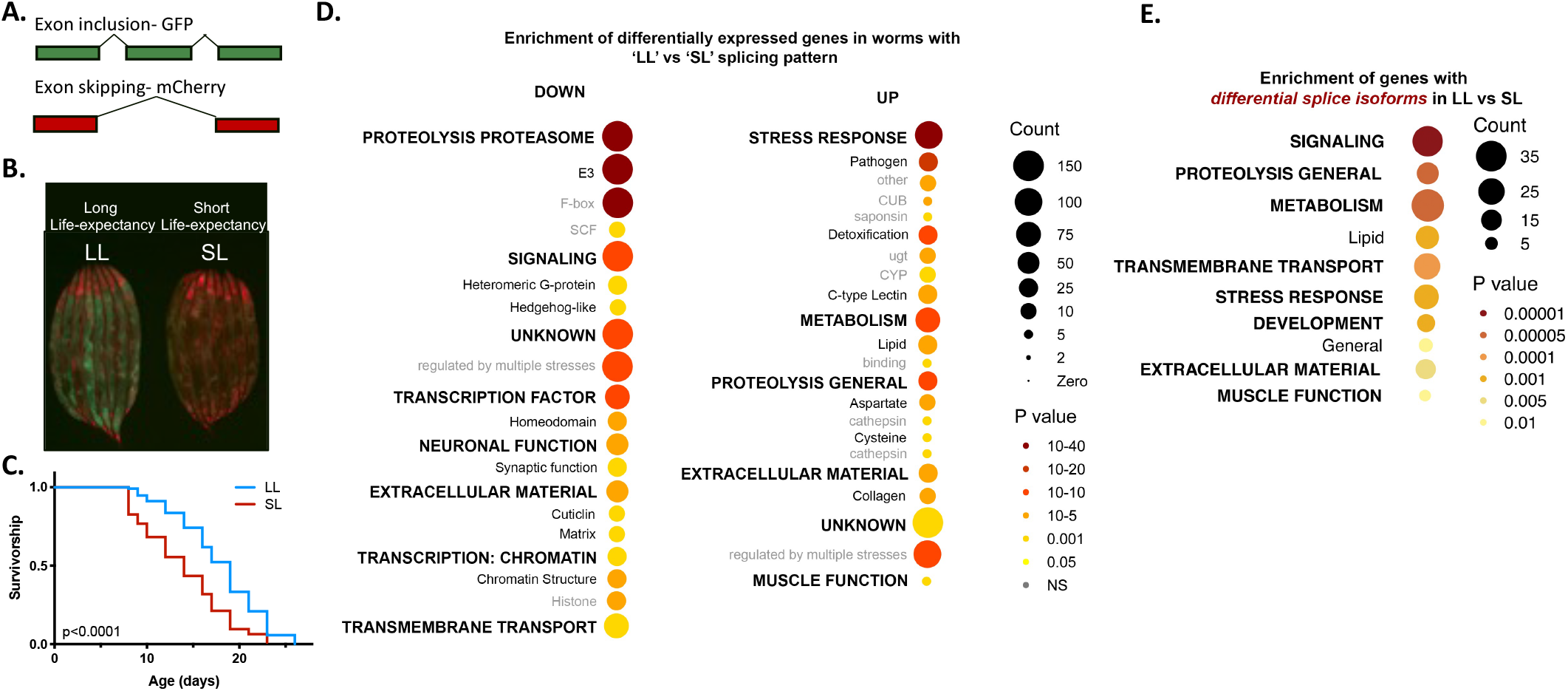
Early life alternative splicing of mRNAs related to lipid metabolism and known longevity pathway correlate with subsequent life expectancy. A. Schematic illustrating the fluorescence expression pattern of *ret-1* splicing reporter worm. Inclusion/exclusion of exon 5 results in GFP/mCherry expression respectively. B. Representative images of *ad-libitum* fed *ret-1* reporter worms segregated at Day 6 based on their splicing pattern. C. Survivorship of the worm sub-populations separated at Day 6 (*P*<0.0001, 1 of 6 replicates). Worms that age slowly are marked ‘LL’ (Long Life-expectancy) and worms that age rapidly as ‘SL’ (Short Life-expectancy). D. WormCat visualization of categories enriched in down and upregulated genes in LL vs SL worms. padj<0.01 and fold change>1.5 was used as cutoff to mark differentially expressed genes. E. Visualization of categories enriched in genes that exhibit differential isoform usage in LL vs SL worms using WormCat gene annotations.

First, we used WormCat (*10*) to identify functional signatures in genes significantly up or downregulated in the long life-expectancy animals (Fig. 1D). LL animals have decreased expression of multiple functional categories including genes regulating signaling and the proteosome, while genes required for stress response/detoxification and proteolysis, processes known to be longevity-related, are increased. LL animals also show widespread changes in genes required for lipid metabolism, with subsets of lipid metabolic genes showing increased expression, suggesting differences in their lipid metabolism or composition (Fig. S1B-E). Differential isoform usage between LL and SL animals is enriched for pathways implicated in aging such as signaling, proteolysis and lipid metabolism further suggesting that splicing of pre-mRNAs specifically linked to these pathways is coupled to life expectancy (Fig. 1E). Notably, differential transcript usage in SL vs LL groups includes genes specific to lipid metabolism, such as *cpt-5, cpt-6, acox-3* and *lpb-1*. Instead of a generalized loss of splicing fidelity in animals that are aging poorly, these data suggest that heterogeneity in expression and splicing of genes in specific functional categories, including lipid metabolism and proteolysis, early in the life of an individual is coupled to life expectancy.

If heterogeneity in RNA splicing of specific functional classes of pre-mRNAs is linked to life expectancy, we reasoned that specific RNA splicing mediated processes may also underlie variation in response to longevity interventions. Previously, we identified causal roles for the RNA splicing factors SFA-1 and REPO-1 in lifespan extension by dietary restriction and suppression of the target of rapamycin complex 1 (TORC1) in *C. elegans* (*6*). REPO-1 (Reversed Polarity-1) is a component of the SF3A subunit in the U2snRNP of the spliceosome whereas SFA-1/ Splicing Factor 1 is a branch-point binding protein that interacts with the UAF proteins and recruits U2 snRNP in the 3’ splice site recognition during early spliceosome assembly (*11*). The link between DR and RNA splicing appears conserved, as upregulation of spliceosome components including SF3A in the liver is a signature of DR in rhesus monkeys (*12*). We sought to define how these RNA splicing factors causally modulate aging, and whether splicing status in an individual might impact its responsiveness to various pro-longevity interventions.

To define whether REPO-1 and SFA-1 activity modulates the efficacy of all longevity interventions or shows specificity, we designed RNAi clones that inhibit REPO-1 or SFA-1 in *C. elegans* without impacting neighboring gene expression (Fig S2). Using these specific RNAi clones, we examined the role of REPO-1 and SFA-1 in a series of long-lived *C. elegans* mutants, targeting distinct longevity pathways. RNAi of REPO-1 and SFA-1 fully suppresses lifespan extension via the DR mimic *eat-2(ad1116)* (Fig. 2A,B), and mutations to multiple components of the TORC1 pathway (*raga-1(ok386), rsks-1(ok1255)*) (Fig. 2C,D, Fig S3A)(*6*) and electron transport machinery (*clk-1(qm30), isp-1(qm150), nuo-6(qm200)*) (Fig. 2E,F; S3B-D). However, knockdown of REPO-1 or SFA-1 had no effect on lifespan extension via mutations to multiple components of the insulin/insulin like growth factor signaling (IIS) pathway (*age-1(hx546)/*PI3K, *daf-2(e1370)/*InR) (Fig. 2G,H, Fig S3E) (*6*), despite blocking lifespan extension by overexpression of the IIS mediator DAF-16/FOXO (Fig. S3F,G). We confirmed *repo-1* RNAi inhibited *repo-1* expression equally in both longevity responsive and non-responsive mutations (Fig. S3H). REPO-1 and SFA-1 activity thus determines responsiveness to specific longevity interventions, rather than modulating global changes that render animals refractory to lifespan extension by any means.

**Figure 2:**
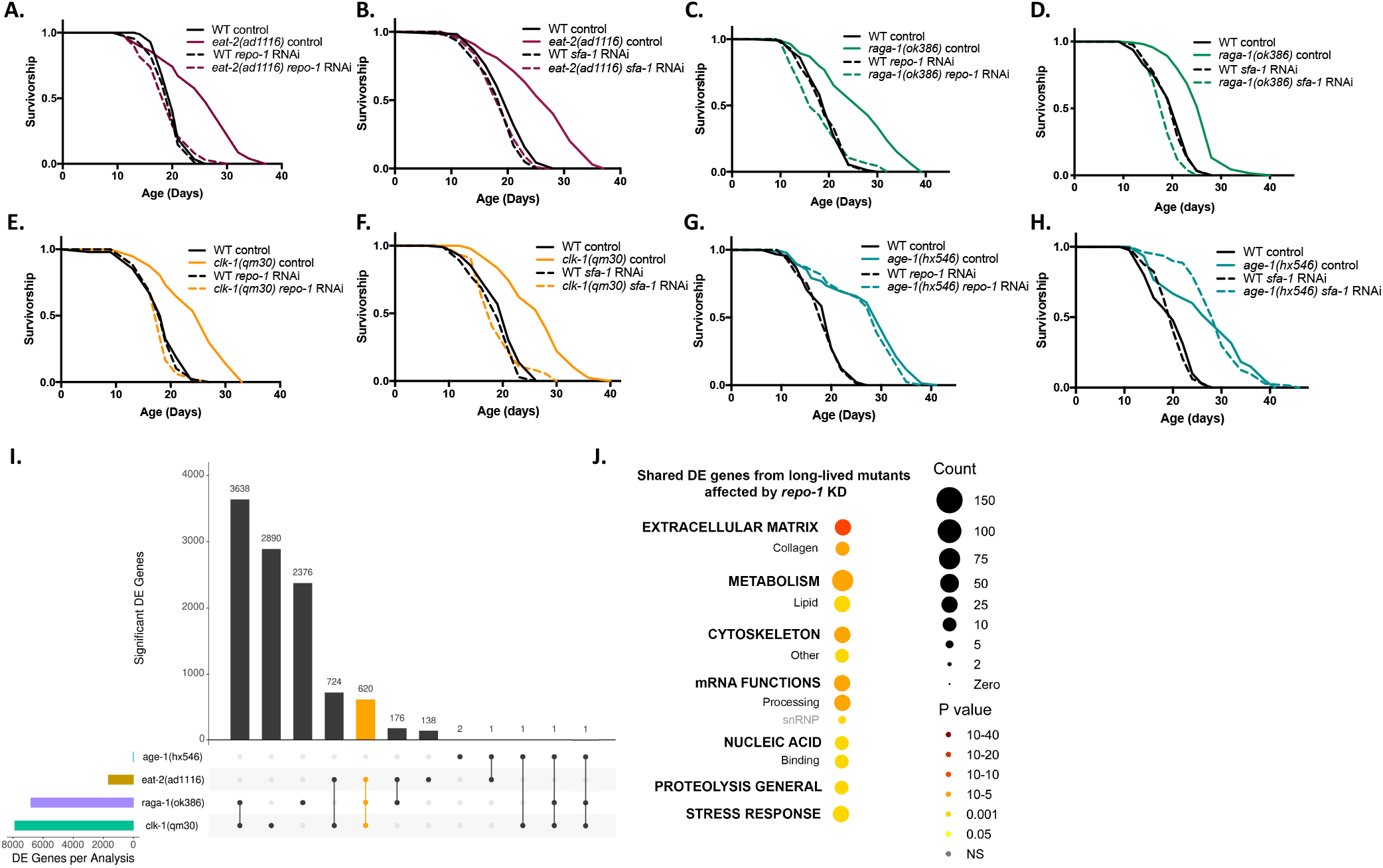
REPO-1 and SFA-1 are required for lifespan extension in DR, TORC1 and ETC mutant longevity but dispensable for IIS longevity. A,B. Survivorship of wild-type (WT) and *eat-2(ad1116)* worms -/+ *repo-1* RNAi (*P*=0.8753) and *sfa-1* RNAi (*P*=0.5658) (*p-*values comparing wildtype N2 on RNAi versus *eat-2(ad1116)* on RNAi, ≥3 replicates). C,D. Survivorship of WT and *raga-1(ok386)* worms -/+ *repo-1* RNAi (*P*=0.5407) and *sfa-1* RNAi (*P*=0.002) (*p-*values comparing wildtype N2 on RNAi versus *raga-1(ok386)* on RNAi, ≥3 replicates). E,F. Survivorship of wild-type (WT) and *clk-1(qm30)* worms -/+ *repo-1* RNAi (*P* = 0.1838 and *sfa-1* RNAi (*P*=0.86) (*p-*values comparing wildtype N2 on RNAi versus *clk-1(qm30)* on RNAi, >=3 replicates). G,H. Survivorship curve of wild-type (WT) and *age-1(hx546)* worms -/+ *repo-1* RNAi (*P*<0.0001) and *sfa-1* RNAi (*P*<0.0001) (*p-*values comparing wildtype N2 on RNAi versus *age-1(hx546)* on RNAi, >=3 replicates). I. UpSet plot (UpSetR R package) quantifying genes in different longevity mutants that respond differently on *repo-1* knockdown compared to wildtype N2 worms. J. WormCat visualization of categories enriched in 620 shared genes that are differentially affected on loss of REPO-1 in splicing factor-dependent pathways compared to wildtype N2 worms.

We leveraged this longevity intervention specificity to define the mechanisms underpinning the effects of REPO-1 on lifespan extension. We examined the transcriptional effects of *repo-1* RNAi on WT animals and a panel of long-lived mutants that either require REPO-1 and SFA-1 for their effects (splicing factor ‘SF’-dependent) or are refractory to them (SF-independent). We performed four replicates of 75-bp paired-end RNA-Seq on wild type (N2), three SF-dependent mutants, *eat-2(ad1116), raga-1(ok386), clk-1(qm30)*, and one SF-independent mutant, *age-1(hx546)*, with and without *repo-1* RNAi. Principal component analysis showed that the transcriptomes of WT, *age-1(hx546)* and *raga-1(ok386)* cluster most tightly, suggesting they share more similarity to each other than to *eat-2(ad1116)* and *clk-1(qm30)* (Fig. S3I). Interestingly, *repo-1* RNAi drove significant changes specifically in *eat-2(ad1116), raga-1(ok386)* and *clk-1(qm30)* mutants yet had a less pronounced effect on WT and *age-1(hx546)* worms, closely mirroring the specific longevity effects of *repo-1* knockdown (Fig. S3I). To identify those changes in gene expression resulting from loss of *repo-1* that are specific to each long-lived mutant, we compared the transcriptional effects of *repo-1* knockdown on each long-lived mutant to the effects of *repo-1* knockdown on wild-type worms. We identified genes whose expression responded differently to *repo-1* RNAi in the long-lived mutants compared to WT. These genes were further categorized and grouped based on the mutant in which they were found (Fig. 2I). Interestingly, in contrast to the widespread differences induced by *repo-1* RNAi in each of the SF-dependent mutants, only 6 genes responded differently on *repo-1* knockdown in the SF-independent mutant *age-1(hx546)* compared to WT worms (Fig. 2J). Together these data suggests that REPO-1 has unique functional roles in the physiology of SF-dependent mutants not seen in WT or IIS mutants, further supporting the idea that REPO-1 activity mediates responsiveness to specific longevity interventions.

We sought to identify the unique role of REPO-1 in SF-dependent longevity interventions. Inhibition of *repo-1* did not induce changes in expression of canonical longevity regulators of the SF-dependent mutants, including the mitochondrial stress response pathway, beta oxidation induction or oxidative stress responses (Fig. S3J-N). We reasoned that processes by which REPO-1 mediates longevity might be reflected in the 620 genes that are differentially regulated by *repo-1* RNAi across all SF-dependent long-lived mutants. We used the WormCat platform (*10*) to identify signatures in these shared 620 differentially regulated genes. REPO-1 dependent gene categories enriched across all splicing sensitive pathways included RNA processing, suggesting indirect or compensatory changes in the post-transcriptional machinery on loss of REPO-1 specifically in these longevity pathways. Strikingly, metabolism was one of the most significantly enriched REPO-1 dependent terms, with lipid metabolism being one of the most enriched sub-categories (Fig. 2J, S4A-D). These data mirror processes coupled to heterogeneity in life expectancy (Fig 1). In addition, we showed previously that loss of SFA-1 specifically reverses expression of lipid metabolism genes induced by DR (*6*). All together these data suggest altered lipid metabolism may be a shared mechanism by which REPO-1 and SFA-1 modulate the aging process and response to treatment.

Next, we examined directly how lipid levels are impacted by *repo-1* and *sfa-1* RNAi in WT and long-lived mutants. Stimulated Raman scattering (SRS) microscopy in live animals showed that neither *sfa-1* and *repo-1* RNAi had any effect on lipid content in the intestine of WT *C*. elegans, yet significantly increased lipid levels in the SF-dependent mutants *eat-2(ad1116)* and *raga-1(ok386)* mutants (Fig. 3A,B). Lipid levels were not affected by loss of REPO-1 or SFA-1 in the splicing resistant mutant *age-1(hx546)*. Interestingly, *sfa-1* and *repo-1* RNAi also had no effect on the lipid content of the splicing sensitive *clk-1(qm30)* mutant, which have impaired mitochondrial function (Fig. 3B), suggesting the mechanism by which these splicing factors impact lipid levels might be linked to mitochondrial function. Therefore, activity of these splicing factor not only specifically regulates expression and splicing of lipid metabolic genes in SF-dependent interventions but lipid content itself.

**Figure 3:**
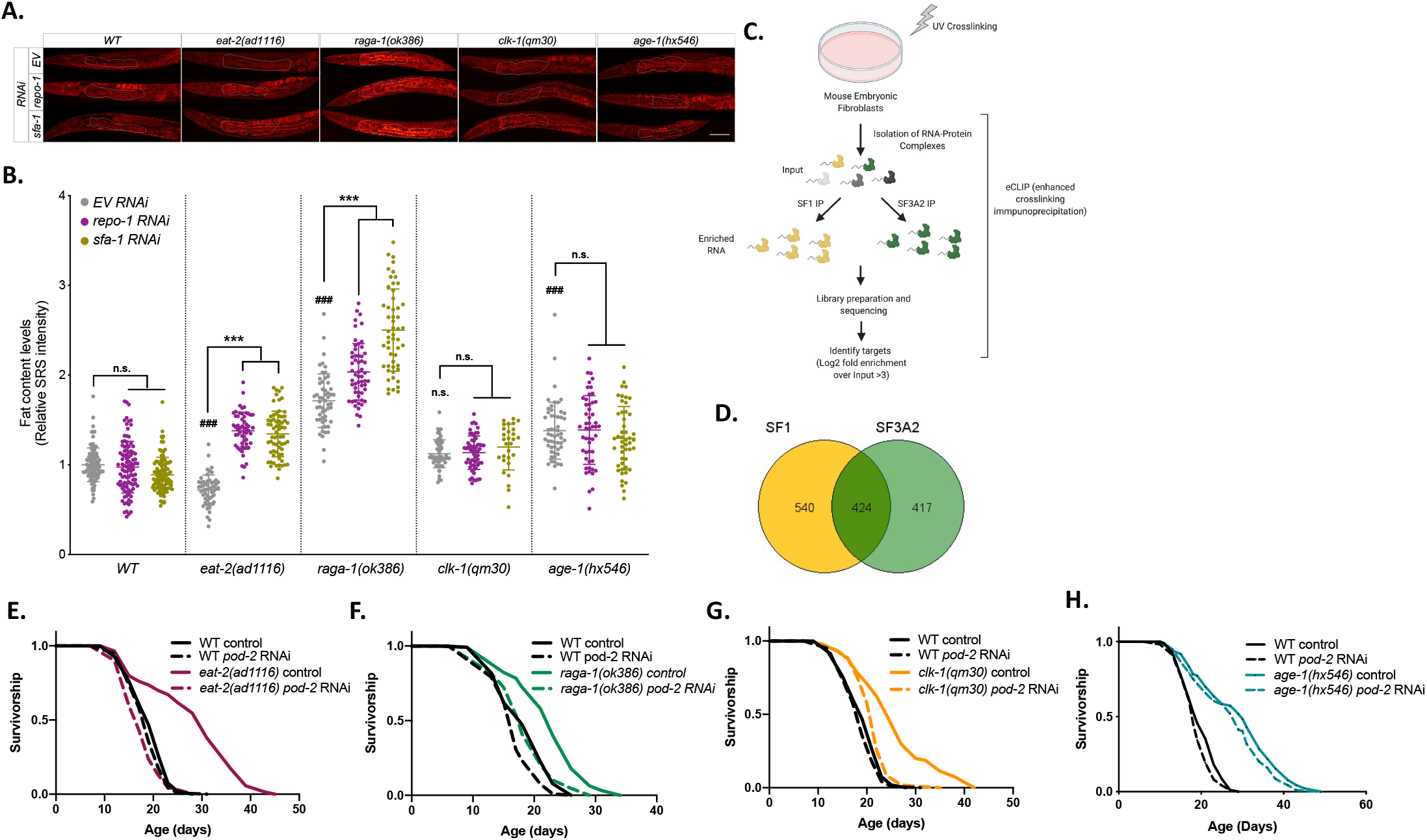
Loss of REPO-1 specifically affects lipid metabolism in splicing factor sensitive longevity pathways. A. Representative images of SRS microscopy showing fat levels in wildtype N2, *eat-2(ad1116), raga-1(ok386), clk-1(qm30)* and *age-1(hx546)* on control, *repo-1* and *sfa-1* RNAi. Worms were fed on RNAi from hatch and imaged 24 hours post L4 stage. B. Quantification of SRS signal using ImageJ (pooled data quantifying anterior intestine of n=20-30 worms, N=3; ### *P*::0.001 long-lived mutant vs wildtype on control EV RNAi; *** *P*::0.001 control EV vs *repo-1/sfa-1* RNAi; n.s. *P*>0.05). C. Schematic of Enhanced Crosslinking Immunoprecipitation (eCLIP) in Mouse Embryonic Fibroblasts (MEFs). D. Venn diagram showing targets of SF1 (mammalian SFA-1 ortholog) and SF3A2 (mammalian REPO-1 ortholog) and their overlap in MEFs. True gene targets identified as peaks in IP over input sample that had a log2fold enrichment>2 and -log10(p-value)>2, N=2. Plot generated using GeneVenn. E-H. Survivorship curves -/+ *pod-2* RNAi of wildtype (WT) and E. *eat-2(ad1116)* (*P*=0.057, 2 replicates); F. *raga-1(ok386)* (*P*=0.1923, 4 replicates); G. *clk-1(qm30)* (*P*<0.0001, 2 replicates) and H. *age-1(hx546)* (*P*<0.0001, 3 replicates). RNAi initiated at Day 1 of adulthood. p-values reflect comparison of wildtype N2 fed with *pod-2* RNAi versus long-lived mutant with *pod-2* RNAi.

To understand the mechanism by which SFA-1 and REPO-1 modulate lipid remodeling and longevity, we identified their direct pre-mRNA targets by enhanced cross-linking immunoprecipitation (eCLIP) (*13*). We performed eCLIP in both *C. elegans* and mouse embryonic fibroblast cell lines (MEFs), speculating that true targets that impact lifespan would be shared between the two splicing factors and conserved across organisms. MEFs were cross-linked followed by IP with SF1 (mammalian SFA-1) and SF3A2 (mammalian REPO-1) antibodies (Fig. 32C). True peaks were defined as those showing enrichment of log2 fold change >= 2 and - log10(p-value) >= 2 over input. Sequencing identified ∼964 and ∼841 genes targeted by SF1 and SF3A2 in MEFs, respectively. 424 genes, representing ∼50% of total targets, are shared, suggesting that SF1 and SF3A2 act together to regulate splicing (Fig. 3D). eCLIP in *C. elegans* shows that many targets of REPO-1 and SFA-1 are also shared in worms, including *tos-1* (Target of Splicing 1), a known and direct target of SFA-1 (*14, 15*) (Fig. S4E-G). Interestingly, in both MEFs and in *C. elegans* we find an enrichment of shared target genes involved in RNA processing and in lipid metabolism. One such target involved in lipid metabolism is Acetyl CoA Carboxylase 1 (ACC1), the rate-limiting enzyme of the fatty acid biosynthetic pathway. ACC1 is a target of both SF1 and SF3A2 in MEFs, and the pre-mRNA for POD-2, the *C. elegans* orthologue of ACC1, was identified as a shared target of SFA-1 and REPO-1 in *C. elegans*.

POD-2/ACC1 converts acetyl-CoA to malonyl CoA, thus catalyzing the first step in the formation of *de novo* lipids. We therefore reasoned that dysfunctional POD-2/ACC1 would alter lipid stores in SFA-1 and REPO-1 depleted animals, and that this might modulate the aging effects in SF-sensitive pro-longevity mutants. To test this, we inhibited lifespan extension of SF-dependent and SF-independent mutants with and without RNAi of ACC/POD-2. Strikingly, inhibition of POD-2 mimicked the longevity intervention specificity of SFA-1 and REPO-1. *pod-2* RNAi from day 1 of adulthood fully suppresses lifespan extension of *eat-2(ad1116), raga-1(ok386)*, and *clk-1(qm30)* (Fig. 3E-G) mutants but does not suppress lifespan in the SF-resistant *age-1(hx546)* (Fig. 3H) mutants. These data suggest that SFA-1 and REPO-1 modulate responses to specific longevity interventions via lipid remodeling and disruption of *de novo* lipid biosynthesis.

Disruption of mitochondrial ETC function and DR/TORC1 inhibition operate in temporally distinct time windows to slow aging in *C. elegans*. Knockdown of ETC complexes only during the L3-L4 stages of larval development increases longevity, while DR and TORC1 inhibition confer longevity benefits in adulthood (*16, 17*). To understand how REPO-1 and SFA-1 modulate longevity and lipid metabolism across seemingly mechanistically unrelated longevity interventions, we asked when in life these splicing factors act to modulate lifespan. We fluorescently tagged endogenous REPO-1 and SFA-1 by CRISPR knock-in of wrmScarlet and GFP respectively. Both splicing factors are expressed in nuclei of all cells (Fig. S5A), throughout all life stages (Fig. S5B) and ages of *C. elegans* (Fig. S5C-E). Neither *repo-1* mRNA nor REPO-1 total protein diminish with age during adulthood (Fig. S5C,D). However, REPO-1 levels peak during early in development, hinting that splicing factors such as REPO-1 play an important role in early life stages of *C. elegans* (Fig. 4A, S5F).

**Figure 4:**
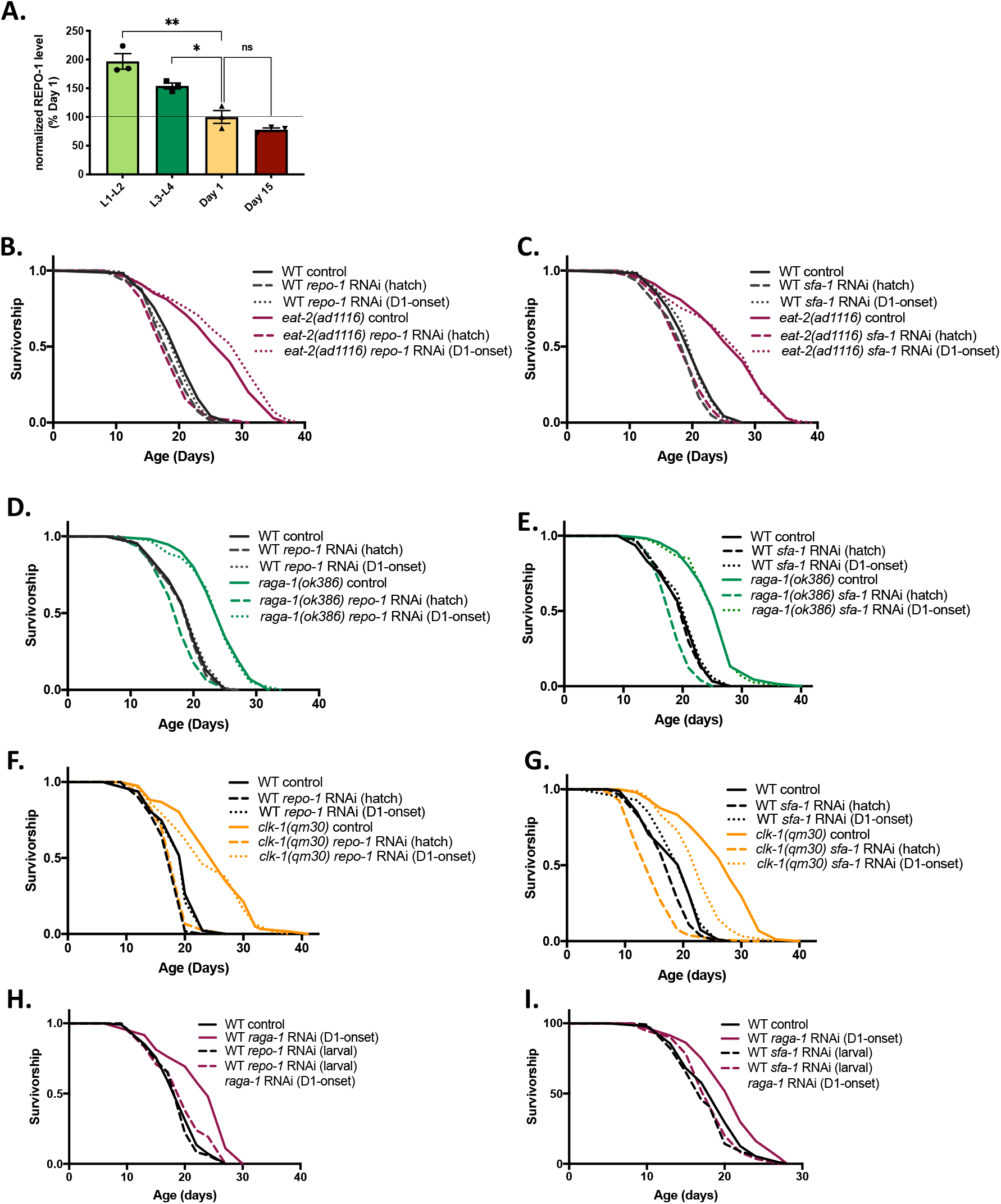
REPO-1 and SFA-1 creates a permissive landscape during the early stages of development to mediate longevity benefits later in life. A. Quantification of bands from western blot of CRISPR-tagged endogenous 3XFLAG::REPO-1 worms at early (L1-L2) and late (L3-L4) larval stages and at Day1 and Day 15 of adulthood. Blots probed with 3XFLAG and actin antibodies. Quantification done using ImageJ, normalized to intensity of actin band and plotted as percent of expression at Day 1 of adulthood (**** *P*≤0.0001, *** *P*≤0.001, ** *P*≤0.01, * *P*≤0.05; ns P>0.05; n=3). B-G. Survivorship of wild-type (WT) and long-lived mutants B. *eat-2(ad1116)*; D. *raga-1(ok386)* and F. *clk-1(qm30)* -/+ *repo-1* from hatch (- - -) or Day 1 of adulthood (D1-onset) (…). P = 0.5315, 0.0285, 0.2672 for WT *repo-1* RNAi hatch vs *eat-2(da1116), raga-1(ok386)* and *clk-1(qm30)* mutants on *repo-1* RNAi hatch respectively). P = <0.0001, <0.0001, <0.0001 for WT *repo-1* RNAi D1-onset vs *eat-2(da1116), raga-1(ok386)* and *clk-1(qm30)* mutants on *repo-1* RNAi D1-onset respectively). Survivorship of wild-type (WT) and long-lived mutants C. *eat-2(ad1116)*; E. *raga-1(ok386)* and G. *clk-1(qm30)* -/+ *sfa-1* from hatch (- - -) or Day 1 of adulthood (D1-onset) (…). P = 0.5658, 0.002, <0.0001 for WT *sfa-1* RNAi hatch vs *eat-2(da1116), raga-1(ok386)* and *clk-1(qm30)* mutants on *sfa-1* RNAi hatch respectively). P = <0.0001, <0.0001, <0.0001 for WT *sfa-1* RNAi D1-onset vs *eat-2(da1116), raga-1(ok386)* and *clk-1(qm30)* mutants on *sfa-1* RNAi D1-onset respectively). G. Survivorship curve of wildtype N2 worms -/+ *repo-1* RNAi in the larval stages -/+ *raga-1* RNAi from Day 1 of adulthood (*P*= 0.0244 wildtype N2 on *repo-1* RNAi (larval) versus wildtype N2 on *repo-1* RNAi (larval)+ *raga-1* RNAi (D1-onset), 3 replicates). H. Survivorship curve of wildtype N2 worms -/+ *sfa-1* RNAi in the larval stages -/+ *raga-1* RNAi from Day 1 of adulthood (*P*= 0.2480 wildtype N2 on *sfa-1* RNAi (larval) versus wildtype N2 on *sfa-1* RNAi (larval)+ *raga-1* RNAi (D1-onset), 2 replicates).

To define the temporal window in which REPO-1 and SFA-1 mediate DR, TORC1 and ETC longevity, we subjected *eat-2(ad1116), raga-1(ok386)* and *clk-1(qm30)* worms to *repo-1* and *sfa-1* RNAi from hatch or Day 1 of adulthood (D1-onset). Despite western analysis confirming equal knockdown efficiency by day 3 of adulthood in both conditions (Fig. S5G-J), their effect on longevity was strikingly different; knockdown of both *repo-1* and *sfa-1* in development fully suppresses longevity in all SF-sensitive mutants. However, adult-onset inhibition had no impact on aging regardless of the timing requirement of the SF-dependent intervention itself (Fig. 4B-G). Together, these data suggest REPO-1 and SFA-1 modulate lipid metabolism early in life and that this permissive landscape can alter effects of subsequent longevity interventions. Supporting this concept, adult-onset RNAi of *raga-1* is sufficient to prolong lifespan yet has no effect on animals with prior inhibition of either *repo-1* or *sfa-1* during development (Fig. 4H,I).

## DISCUSSION

Geroscience approaches over the last 20 years have shown that environmental conditions and genetic manipulation can strongly influence the rate of physiological aging. In particular, DR has a strong beneficial effect in a wide variety of organisms tested to date, not only increasing longevity but also protecting against many chronic diseases. Harnessing the molecular and cellular processes mediating the plasticity of aging in response to DR has the potential to yield novel therapeutics that broadly reduce disease incidence in the elderly. However, while there has been much research into the molecular mechanisms mediating longevity interventions using model organisms, little emphasis has been placed on predicting and understanding variance in their efficacy (*18*). This is critical, since the same therapeutic treatment can be beneficial or harmful, depending on the sex, genotype and physiological state of the individual to which it is applied. For example, dosage levels of pharmacological treatments or DR have very different longevity effects in different genotypes (*19, 20*). Such genotype by diet interaction may well explain the opposing results of two longitudinal lifespan studies of DR on rhesus monkeys (*21*).

Beyond inter-genotype variation to longevity interventions, substantial variation also exists in inter-individual responses. Even within isogenic populations of *C. elegans*, or inbred *Drosophila* and mouse strains, high variance exists both for lifespan itself and response to DR/DR mimetics. Indeed, for interventions such as methionine restriction, though median lifespan of mice and rats is increased, a sub population die earlier than non-restricted controls (*22*). A ‘one size fits all’ approach to DR or DR mimetics is therefore unlikely to be useful therapeutically. Successfully translating geroscience to human application requires accurately predicting a given individual’s response to a given treatment, depending on the genotype and physiology of the individual. Such an approach will allow treatments to be assigned to each individual in a personalized way, maximizing health benefits. The first step toward this end is to identify biological variables that predict individual-specific optimal aging interventions in model organisms, to provide the foundation for a personalized medicine approach to healthy aging therapeutics in humans.

Here we show that in *C. elegans*, a single RNA exon skipping event can predict life expectancy of isogenic animals. We leveraged this finding to define functional categories of genes that are differentially expressed or spliced in animals that are aging well versus those aging poorly. Animals that naturally age well show gene expression and RNA splicing changes enriched in specific ontology classes, including signaling, proteolysis and lipid metabolism. We utilized splicing factor dependent and independent longevity pathways to define RNA splicing mediated processes coupled to variation in response to anti-aging interventions in *C. elegans*. Regulation of genes tied to lipid metabolism was a shared process linking both WT life expectancy and response to treatment. REPO-1 and SFA-1 activity modulate lipid levels in SF-dependent mutants and bind to POD-2/ACC directly. Suggesting a causal link between lipid remodeling and response to treatment, REPO-1, SFA-1, and POD-2 specifically modulate efficacy of the same pro-longevity mutations. Lastly, we show that modulating the activity of REPO-1 and SFA-1 early in life determines whether subsequent application of a pro-longevity intervention, in this case inhibition of RAGA-1, successfully slows aging.

These data are proof of principle that the efficacy of a pro-longevity intervention is strongly influenced by prior events, in this case the activity of a splicing factor and lipid metabolic landscape earlier in the life of the individual. If conserved in mammals, this raises important questions about how we design and implement anti-aging therapeutics. Ultimately it might be possible to predict response to treatment and optimize the intervention via analysis of a subset of RNA splicing events. Such an approach is key if we are to move beyond biomarkers of physiological age towards biomarkers that facilitate precision medicine approaches to interventional geroscience.

## Acknowledgements

We are thankful to H. Kuroyanagi for sharing the *ret-1* splicing reporter strain. We thank *Caenorhabditis Genetics Center* for providing worm strains. We are also grateful to members of the Mair laboratory for helpful discussions on the project and the manuscript.

## Funding

WBM is funded by the National Institutes of Health, NIA/NIH RO1 R01AG051954, NIA/NIH RO1 AG059595, NIA/NIH R01AG067106, NIA/NIH R01AG054201 and NIA/NIH R01AG044346. This work was also supported by the American Federation for Aging Research/ Glenn Foundation for Medical Research Breakthroughs in Gerontology Award. MCW is funded by HHMI and R01AG062257. Work done by MEP at the Harvard Chan Bioinformatics Core was partially subsidized by the Harvard Stem Cell Institute.

## Author Contributions

S.D. and W.B.M. designed the study; S.D. performed most experiments and analyzed results, C.H. performed experiments and edited the manuscript; M.C.P.M., A.S. H.J.S. P.H. and R.S. performed lifespan repeats; M.E.P., M.M. and M.C.P.M performed analysis of the RNA-Sequencing data; A.S.M. performed SRS Microscopy in M.C.W. Lab; S.D., A.L. and W.B.M. wrote the manuscript, implemented comments and edits from all authors.

## Competing Interests

Authors declare no competing interests.

## Data and Materials Availability

All data will be deposited to a public database before publication.

## SUPPLEMENTARY MATERIAL

### MATERIALS AND METHODS

#### Worm strains and culture

The *C. elegans* strains used for this work are listed below. Worms were routinely grown and maintained on standard nematode growth media (NGM) plates that were seeded with *E. coli* (OP50-1). *E. coli* bacteria were cultured overnight in LB at 37°C, after which 100 µl of liquid culture was seeded on plates to grow for 2 days at room temperature.

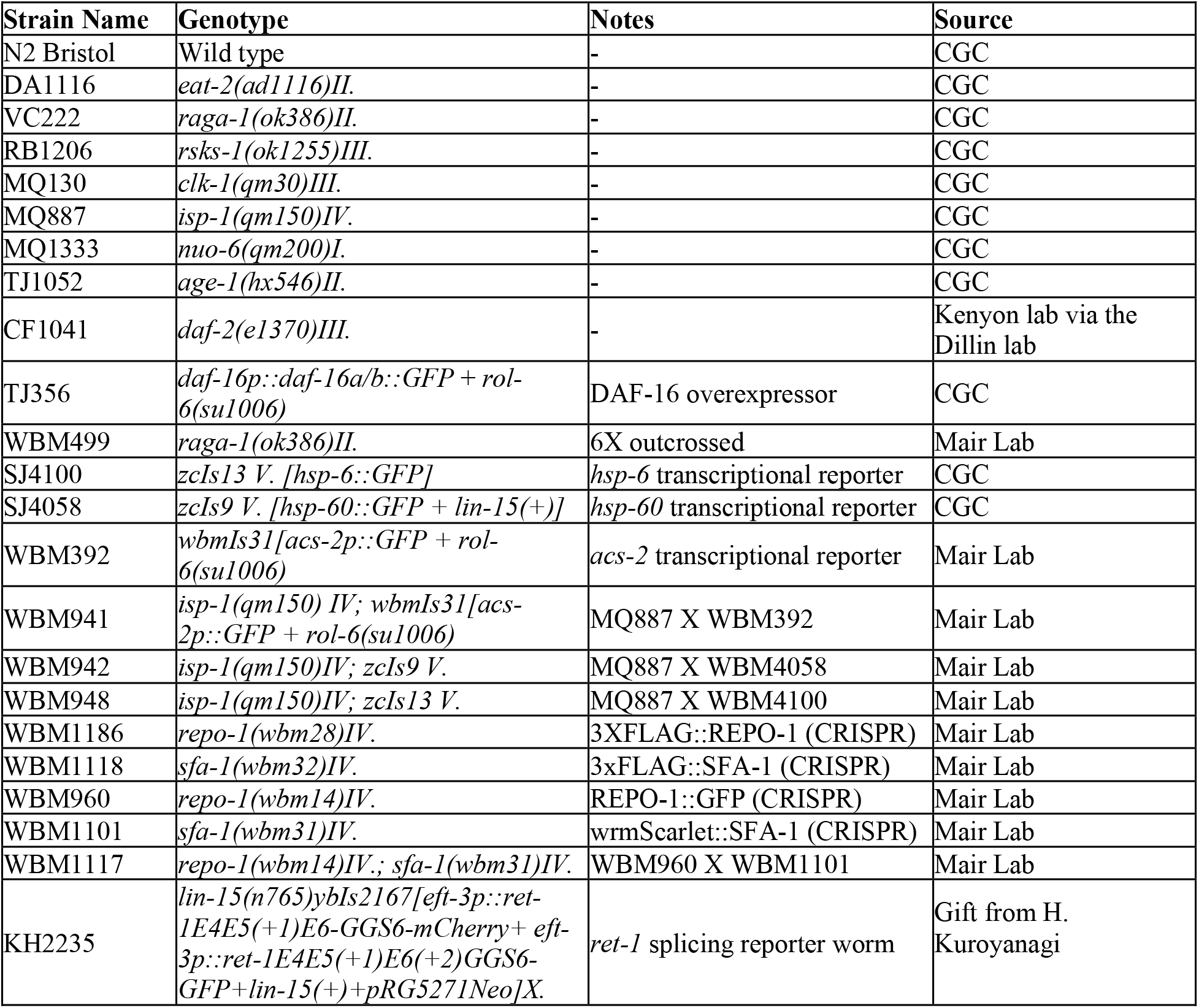

| Strain Name | Genotype | Notes | Source |
| --- | --- | --- | --- |
| N2 Bristol | Wild type | - | CGC |
| DA1116 | *eat-2(ad1116)II.* | - | CGC |
| VC222 | *raga-1(ok386)II.* | - | CGC |
| RB1206 | *rsks-1(ok1255)III.* | - | CGC |
| MQ130 | *clk-1(qm30)III.* | - | CGC |
| MQ887 | *isp-1(qm150)IV.* | - | CGC |
| MQ1333 | *nuo-6(qm200)I.* | - | CGC |
| TJ1052 | *age-1(hx546)II.* | - | CGC |
| CF1041 | *daf-2(e1370)III.* | - | Kenyon lab via the Dillin lab |
| TJ356 | *daf-16p::daf-16a/b::GFP + rol-6(su1006)* | DAF-16 overexpressor | CGC |
| WBM499 | *raga-1(ok386)II.* | 6X outcrossed | Mair Lab |
| SJ4100 | *zcls13 V. [hsp-6::GFP]* | *hsp-6* transcriptional reporter | CGC |
| SJ4058 | *zcls9 V. [hsp-60::GFP + lin-15(+)]* | *hsp-60* transcriptional reporter | CGC |
| WBM392 | *wbmls31[acs-2p::GFP + rol-6(su1006)]* | *acs-2* transcriptional reporter | Mair Lab |
| WBM941 | *isp-1(qm150) IV; wbmls31[acs-2p::GFP + rol-6(su1006)]* | MQ887 X WBM392 | Mair Lab |
| WBM942 | *isp-1(qm150)IV; zcls9 V.* | MQ887 X WBM4058 | Mair Lab |
| WBM948 | *isp-1(qm150)IV; zcls13 V.* | MQ887 X WBM4100 | Mair Lab |
| WBM1186 | *repo-1(wbm28)IV.* | 3XFLAG::REPO-1 (CRISPR) | Mair Lab |
| WBM1118 | *sfa-1(wbm32)IV.* | 3xFLAG::SFA-1 (CRISPR) | Mair Lab |
| WBM960 | *repo-1(wbm14)IV.* | REPO-1::GFP (CRISPR) | Mair Lab |
| WBM1101 | *sfa-1(wbm31)IV.* | wrmScarlet::SFA-1 (CRISPR) | Mair Lab |
| WBM1117 | *repo-1(wbm14)IV.; sfa-1(wbm31)IV.* | WBM960 X WBM1101 | Mair Lab |
| KH2235 | *lin-15(n765)ybls2167[eft-3p::ret-1E4E5(+1)E6-GGS6-mCherry+ eft-3p::ret-1E4E5(+1)E6(+2)GGS6-GFP+lin-15(+)+pRG5271Neo]X.* | *ret-1* splicing reporter worm | Gift from H. Kuroyanagi |

#### Lifespans

All lifespans were conducted at 20°C. Lifespans were performed as described in (*1*). Graphpad Prism 9 was used to plot survival curves and calculate median lifespans. Lifespans were started with n=100 or 120 worms for each condition unless otherwise specified. Survival curves were compared, and p-values were calculated using the log-rank (Mantel-Cox) analysis method.

#### RNA interference

RNAi construct for *sfa-1* was obtained from the Ahringer RNAi and sequence verified (*2*). The *repo-1* RNAi was made by cloning the first 403 bases of the *repo-1* gene into the L4440 vector. RNAi experiments were done using *E*.*coli* HT115 bacteria on standard NGM plates containing 100µg/ml Carbenicillin. HT115 bacteria expressing RNAi constructs were grown overnight in LB supplemented with 12.5µg/ml Tetracycline and 100µg/ml Carbenicillin. The plates were seeded with 100 µl of the bacterial culture 48 hours before use. Respective dsRNA expressing HT115 bacteria were induced by adding 100µl IPTG (100mM) 1-2 hours before introducing worms to the plate. RNAi was induced from hatch or at Day 1 of adulthood as specified. Worms grown on empty vector HT115 RNAi bacteria are represented as control.

The sequences of the different RNAi constructs are listed below:

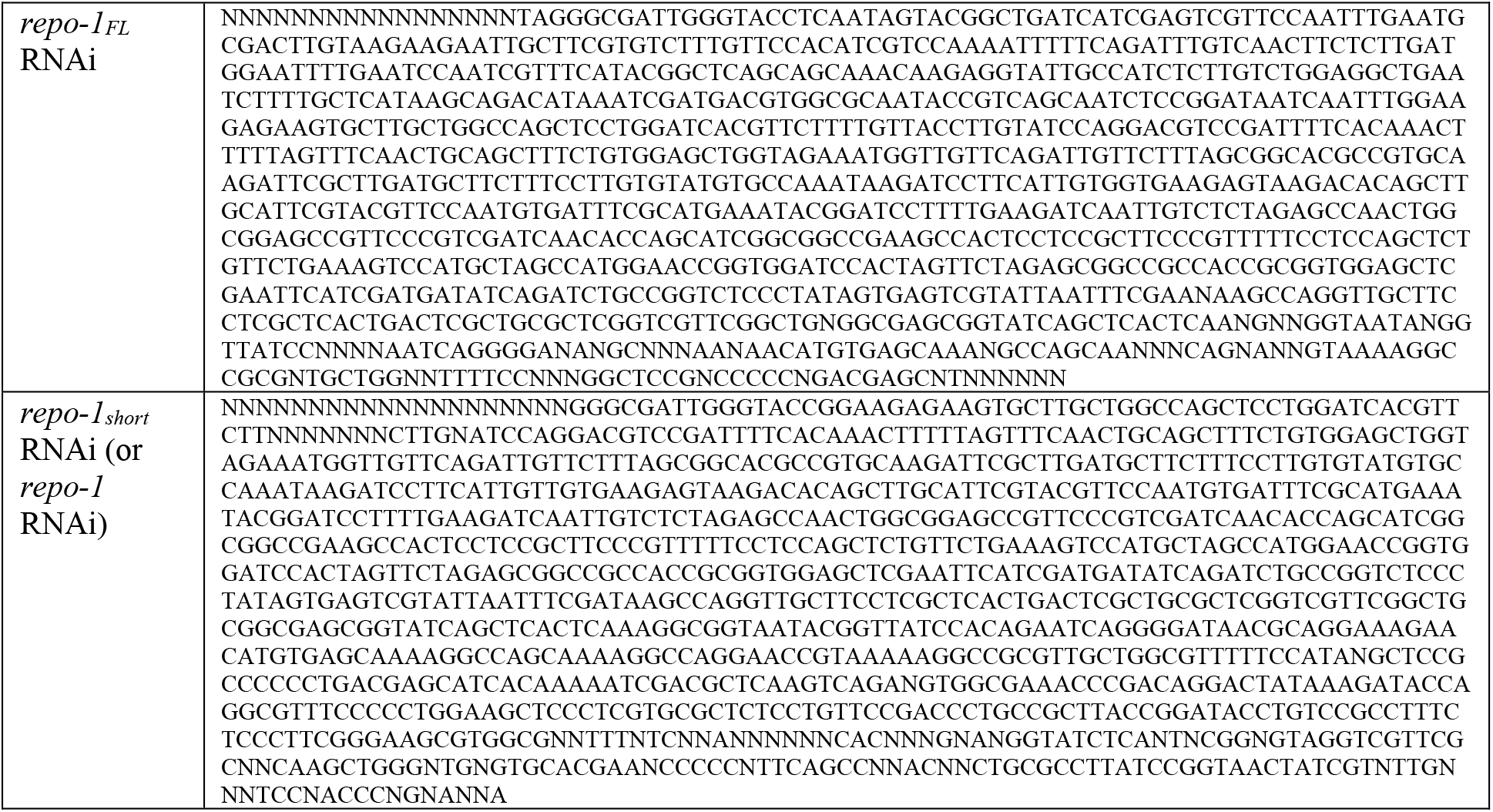
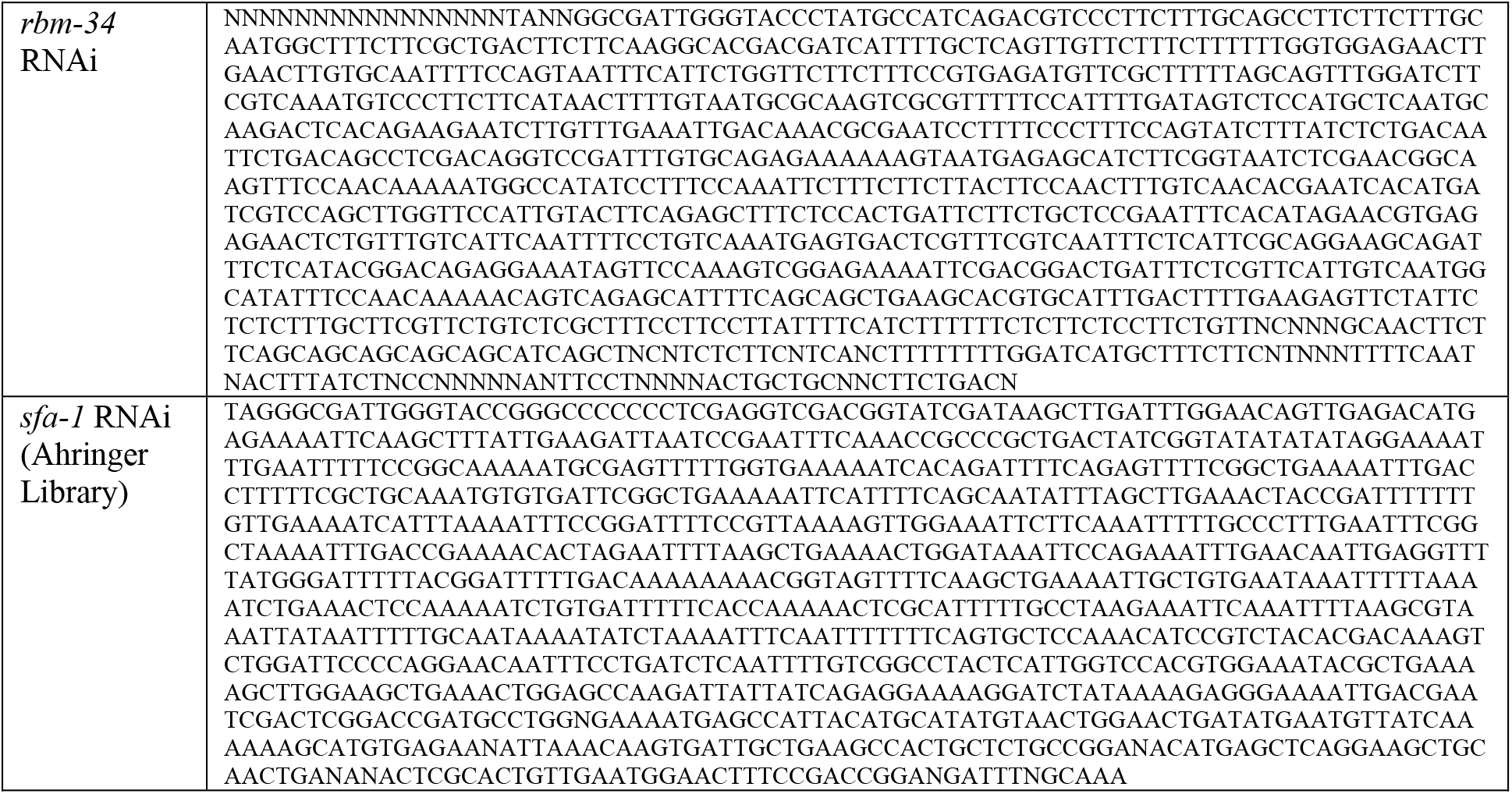

#### Sorting of splicing reporter worms

KH2235 splicing reporter worms were thawed from -80°C stocks for each independent biological replicate. Worm populations were grown on NGM plates seeded with *E. coli* OP50-1 and age synchronized using 5% sodium hypochlorite solution. Around 2500 *C. elegans* eggs were placed on NGM plates with OP50-1 bacteria following the bleach process. On the first four days of adulthood, worms were transferred daily to fresh plates to remove progeny. At day 6 of adulthood, worm populations were assigned by visual separation to respective youthful and aged splicing pattern groups using a UV dissecting microscope (Zeiss Discovery V8). From each group separately, 100 were randomly selected using white light to include in survival analysis. Lifespans were set up as previously described. In addition, 8 randomly selected animals from each group were imaged as a representative population of the group as described previously (*2*).

For RNA extractions, about 200 animals were washed off plates using M9+0.01% Tween, washed twice using M9 buffer and flash-frozen in 250µl Qiazol in liquid nitrogen. For each biological replicate, duplicate samples were frozen. Samples were stored at -80°C and RNA extractions were performed in parallel for all biological replicates.

RNA extractions were performed using Qiagen miRNeasy micro kit and eluted in ddH_2_O. Six independent biological replicates were sent for library preparation and RNA sequencing.

Three independent biological replicates were collected after the RNA sequencing following the same protocols for validation by qRT-PCR.

### RNA isolation and cDNA synthesis

Total RNA was extracted using Qiazol (QIAGEN), column purified by RNeasy mini kit (QIAGEN) according to manufacturer’s instructions. cDNA was synthesized using SuperScript® VILO Master mix (Invitrogen).

#### Quantitative RT-PCR

StepOne Plus instrument from Applied Biosystems was used to perform real-time qPCR experiments following the manufacturer’s instructions. The following Taqman assays from Life Technologies were used: *repo-1* (Ce02465496_g1), *rbm-34* (Ce02465498_g1), *acs-2* (Ce02486192_g1), *sod-3* (Ce02404518_gH), *fat-3* (Ce02458252_g1), *cpt-5* (Ce02419317_g1), *fat-7* (Ce02477067_g1), *acdh-1* (Ce02408341_g1), *acdh-2* (Ce02432818_g1), *gst-10* (Ce02504848_m1) . For each qPCR reaction, ∼5ng of cDNA was used. Relative expression differences were calculated with the comparative 2ΔΔCt method using Y45F10D.4 (Ce02467253_g1) as the endogenous control. For each gene in each strain, fold-change relative to the average of wild type control group was calculated and statistical significance evaluated using Welch’s t test. Graphpad Prism 9 was used for all statistical analysis and graph plotting.

#### Semi-quantitative RT-PCR of *tos-1* to visualize alternative splicing event

*tos-1* was amplified using PCR conditions and primers as described in (*2*). Apex Taq RED (Genesee Scientific) was used for amplification. 1 Kb Plus DNA ladder (Invitrogen) was used as molecular weight reference. Following PCR, samples were resolved on a 2% EtBr-stained agarose gel. Gels were imaged on ChemiDoc MP (BioRad).

#### Microinjection and CRISPR/Cas9 mediated gene editing

All CRISPR edits were performed using the CRISPR protocol developed by (*3*). Briefly, homology repair templates were amplified by PCR using primers that introduced a minimum stretch of 35 bp homology arms flanking the site of insertion at both ends. The CRISPR injection mix contained 2.5 μl tracrRNA (4 μg/μl), 0.6 μl *dpy-10* crRNA (2.6 μg/μl), 0.5 µl target gene crRNA (2.6 μg/μl), 0.25 μl *dpy-10* ssODN (500 ng/μl), homology repair template (200 ng/μl final in the mix), 0.375 μl Hepes pH 7.4 (200mM), 0.25 μl KCl (1M) and RNase free water to make up the volume to 8 µl. 2 μl purified Cas9 (12 μg/μl) was added at the end, mixed by pipetting, spun for 2 min at 13000 rpm and incubated at 37°C for 10 minutes. *dpy-5* was used as a co-injection marker instead of *dpy-10* in case the edits were made on Chromosome II. Mixes were microinjected into the germ line of day 1 adult hermaphrodites using standard protocol. Screening worms and genotyping was performed as described earlier (*4*). Worms generated using CRISPR were outcrossed at least six times before being used for experiments to remove the co–injection marker phenotype and other off-target edits.

#### Microscopy

For DIC and fluorescence imaging of REPO-1::GFP and wrmScarlet::SFA-1 at different stages, worms were anesthetized in 0.1 mg/ml tetramisole in 1X M9 buffer on empty NGM plates and mounted on 2% agarose pads on glass slides. They were imaged using the Apotome.2-equipped Imager M2 microscope with an Axiocam camera. For imaging of hsp-6 and hsp-60, on NGM plates without bacteria until no movement was detectable, aligned to groups accordingly and subsequently imaged on a Zeiss Discovery V8 microscope equipped with an Axiocam camera. Exposure times were kept constant for all imaging experiments involving the splicing reporter.

#### Western Blotting and Quantification

3XFLAG::*repo-1* worms were egg laid on *E. coli* HT115 bacteria to get a synchronized population of worms. To age them up until Day 15, worms were transferred on days 1, 2, 3, 5 and 8 to separate them from their progeny. For Day 1, Day 5, Day 10 and Day 15 samples, approximately 200 adults per sample were collected in triplicates in M9 buffer and snap frozen in liquid nitrogen. For larval stages, approximately 40 animals were made to egg lay over a 12-14 hour period on each plate. 10 such plates were pooled after 24 or 48 hours to get one replicate of L1-L2 or L3-L4 stage sample respectively. To make worm lysates, RIPA buffer with protease inhibitors (Sigma #8340) were added and samples were lysed via sonication at Amplitude 60 10sec ON / 10sec OFF for a total of 3 times (Qsonica Q700) and this complete cycle was repeated 3 times. SDS-PAGE was performed by running 20 µg of protein per lane on a 4-12% TrisGlycine gradient gel (Thermo Fisher Scientific, #XP04122BOX). Proteins were transferred to Nitrocellulose membranes (BioRad # 162-0112) and blocked with 5% BSA in TBST. They were stained with Ponceau-S Stain, 0.1% Solution (G Biosciences #89167-800) and imaged. Ponceau was washed off with 1XTBST and blots were incubated with primary antibody ANTI-FLAG M2 mouse monoclonal (Sigma Aldrich #F1804, 1:2000). They were then washed in 1XTBST and probed with secondary anti-HRP linked mouse antibody (Cell Signaling #7076, 1:5000). Blots were developed using ECL substrate (GE Healthcare # 95038-560). Bands were visualized using a Gel Doc system (Bio Rad) and Image Lab software (Version 4.1). Blots were stripped using Stripping Buffer (Thermofisher # PI46430) according to manufacturer’s guidelines and re-probed with beta actin (Abcam, #8226, 1:1000). Quantification of the bands was done using ImageJ (Version 1.52a) and plotted using Graphpad Prism 9.

#### RNA-Sequencing

Libraries were prepared using Roche Kapa stranded mRNA HyperPrep sample preparation kits from 100ng of purified total RNA according to the manufacturer’s protocol. The finished dsDNA libraries were quantified by Qubit fluorometer, Agilent TapeStation 2200, and RT-qPCR using the Kapa Biosystems library quantification kit according to manufacturer’s protocols. Uniquely indexed libraries were pooled at an equimolar ratio and sequenced on an Illumina NextSeq500 with paired-end 75bp reads by the Dana-Farber Cancer Institute Molecular Biology Core Facilities.

#### Differential Gene Expression Analysis

All samples were processed using an RNA-seq pipeline implemented in the bcbio-nextgen project (https://bcbio-nextgen.readthedocs.org/en/latest/). Raw reads were examined for quality issues using FastQC (v0.11.8) (http://www.bioinformatics.babraham.ac.uk/projects/fastqc/) to ensure library generation and sequencing were suitable for further analysis.

To perform additional quality checks of the data, all reads were aligned to Wormbase assembly WBcel235, release WS272 of the C. elegans genome (Project PRJNA13578 (N2 strain)) using STAR (v. 2.6.1d) (*5*). Alignments were checked for evenness of coverage, rRNA content, genomic context of alignments (for example, alignments in known transcripts and introns), complexity and other quality checks using a combination of FastQC, Qualimap (*6*), http://doi.org/10.1093/bioinformatics/bts503], MultiQC (https://github.com/ewels/MultiQC) and custom tools. For the LL vs SL worm RNA-seq, the reference genome annotation file used was WBcel235, ensemble release v99.

To quantitate the reads corresponding to each transcript, quasi alignment was performed using Salmon (v. 0.14.2) (*7*) in both datasets, outputting the transcripts Per Million (TPM) measurements per isoform. Differential expression at the gene level was called with DESeq2 (*8*), using counts per gene estimated from the Salmon quasi alignments by tximport. The differential effect of *repo-1* knockdown on the mutants *raga-1(ok386), eat-2(ad1116)*, and *clk-1(qm30)* relative to *age-1(hx546)* was explored using the design formula: ∼ repo1_effect + treatment + repo1 effect:treatment.

The DEGReport (*9*) [http://lpantano.github.io/DEGreport/] package was used to identify gene clusters that change similarly upon knockdown of repo-1 using a hierarchical correlation clustering approach. UpSet plots of shared differentially expressed genes between analyses were generated using the UpSetR R package (*10*). Lists of differentially expressed (DE) genes in LL vs SL worm RNA-Seq or groups of DE genes with similar expression changes with *repo-1* knockdown in REPO-1 RNA-Seq identified by DEGReport were examined for enrichment using WormCat (*11*).

#### Splicing and Differential Isoform Usage Analysis of SL vs LL worm sub-populations

Differential transcript usage and local alternative splicing events were identified using SUPPA2 (*12, 13*). The TPM values output from Salmon were used as input to SUPPA2, and differential splicing analysis was performed using the empirical method. The p-values were corrected for multiple testing and an alpha of 0.05 was used for identification of significant events. An event was not tested if none of the transcripts associated with the event were expressed for any sample or if one or more transcripts of the event was not quantified for any sample.

The genes corresponding to significant events were examined for over-representation analysis with clusterProfiler (*14*). The background set of genes represent all spliced genes that were tested for differential splicing of events (e.g. spliced genes that were expressed in all samples). The significant genes were de-duplicated for genes that corresponded to more than one significant event. Enriched processes represent those biological processes enriched for genes that are significantly differentially spliced compared to all, expressed spliced genes. The biological processes tested were defined by the gene sets obtained from Wormcat (*11*) and were tested separately at each hierarchical level.

#### SRS microscopy imaging, *C. elegans* sample preparation and image quantification

For the SRS microscope system, pulsed Pump (tunable from 790 to 990 nm) and Stokes (1045 nm) beams were provided by Insight X3 femtosecond laser (Spectra-Physics), and spatiotemporally overlapped by a Spectral Focus Timing and Recombination Unit (SF-TRU, Newport). The intensity of Stokes beam is modulated at 20 MHz by an electro-optic modulator (EOM, 4103, New Focus). Overlapping pump and Stokes beams were emitted from the port of SF-TRU and coupled into a multi photon laser scanning microscope (FVMPE-RS, Olympus). After passing through the sample, the forward going Pump and Stokes beams were collected by an air condenser. A flip mirror was used to direct the transmitted laser to a photodiode module, which includes a telescope to relay and change the beam size to fit the area of the photodiode, as well as an optical filter to remove the modulated Stokes beam and let the pump beam transmit for the detection of stimulated Raman loss signal. The output current from the photodiode was terminated, filtered, and demodulated by a lock-in amplifier (SRS Detection Module, APE) at 20 MHz to ensure shot noise-limited detection sensitivity. The lock-in amplifier output was then fed into the analog box of FVMPE-RS microscope system to process the SRS signal. For lipid imaging, CH_2_ signals were detected at 2845 cm^-1^. To achieve this, pump beam was set at 805 nm and the delay stage was positioned at 33.550 nm. For both pump and Stokes beams, 10% laser power was used. A 20x air objective (UPlanSAPO, 0.75 N.A., Olympus) was used for imaging. The microscope was controlled by Olympus Fluoview software. SRS microscopy images were quantified using ImageJ software (NIH). Polygon selection tool was used to select the area to be quantified and average pixel intensity was calculated with the “analyze-measure” command. After subtracting the background intensity, all measurements were averaged to obtain mean and standard deviation. In each imaging session, approximately 20-30 worms were immobilized with 1% sodium azide on 2% agarose pads on glass microscope slides and 10-20 worms, which have anterior intestine in focus, were imaged. Mutant lipid levels were normalized to those of wild-type worms and relative SRS signals are shown in box plots. Wild-type and control worms were bleached on plates with HT115 bacteria and their progeny were used for adult-egg laying on plates with HT115 bacteria transformed with either empty L4440 vector or vector carrying RNAi against *sfa-1* or *repo-1*. Embryos were grown at 20°C, on these plates, until larval L4 stage, when they are synchronized again and imaged 24 hours later.

#### Enhanced Crosslinking Immunoprecipitation (eCLIP) in MEFs and Worms

eCLIP was performed by Eclipse Bioinnovations Inc (San Diego, www.eclipsebio.com) according to the published eCLIP protocol (*15*). For each replicate (N=2), 20 million MEFs were UV crosslinked (UVC-515 Ultralum) at 400 mJoules/cm2 with 254-nm radiation, and snap frozen in liquid nitrogen. Cells were lysed and treated with RNase I to fragment RNA as previously described. SF1 (Bethyl: Cat No. A303-213A, Lot No. A303-213A-1) and SF3A2 (Bethyl: Cat No. A304-821A, Lot No. A304-821A-1) antibodies were used for immunoprecipitation of the proteins respectively. Only the region from 65 kDa to 140 kDa was excised for eCLIP. RNA adapter ligation, immunoprecipitation-western blotting, reverse transcription, DNA adapter ligation and PCR amplification were performed as previously described.

For *C. elegans* eCLIP (N=1), 50,000 worms or 50uL packed worms (∼1mg of total protein) was grown. Worms were obtained from an egg lay of CRISPR tagged 3XFLAG::SFA-1 (WBM1118) and 3XFLAG::REPO-1 (WBM1186) on 10 cm plates seeded with HT115 bacteria. At Day 1 of adulthood, they were transferred to a 15 ml centrifuge tube in M9+0.01% Tween and washed gently 3 times with 10mL M9 at room temperature. They were resuspended in 5 ml fresh M9 and transferred to NGM plates without bacteria. After the worms dispersed uniformly, the plates were placed on leveled ice, plate lid removed followed by crosslinking (UVC-515 Ultralum) at 254-nm UV with an energy setting of 500 mJoules/cm2. Immobilized worms were transferred to a 2mL round bottom microcentrifuge tube using M9 and spun for 30 seconds at 3,000xg. Excess M9 was aspirated out and the worm pellet flash frozen in liquid nitrogen. Monoclonal Anti-FLAG M2 antibody (Sigma: Cat No. F1804-1MG, Lot No. SLBX2256) was used for immunoprecipitation of FLAG::SFA-1 and FLAG::REPO-1 and eCLIP was performed by excising the 80-155kDa region for SFA-1 and 32-115kDa for REPO-1.

## SUPPLEMENTARY FIGURE LEGENDS

**Figure S1:**
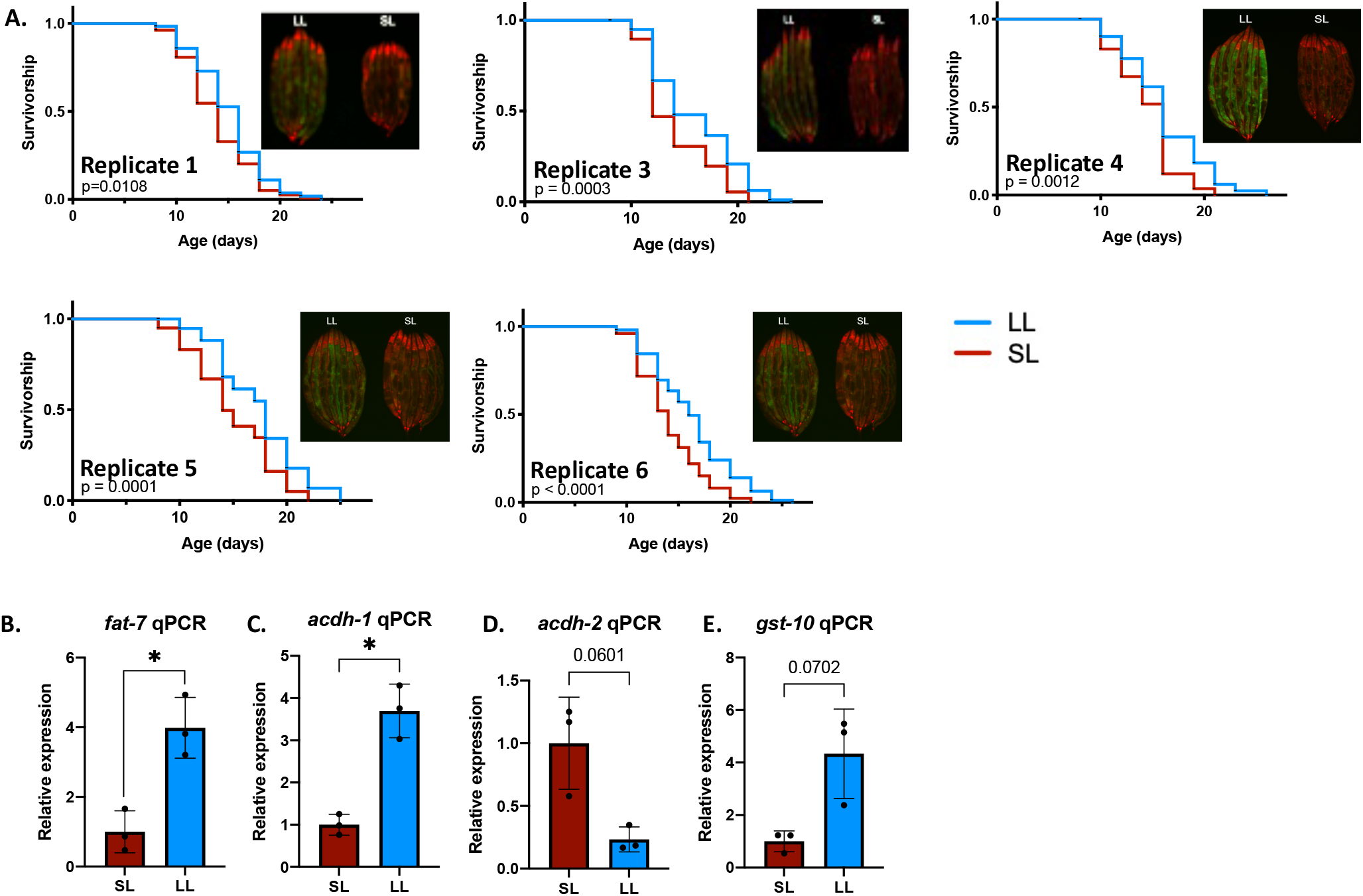
Early life alternative splicing of RNAs related to lipid metabolism and known longevity pathway correlates with subsequent life expectancy. A. Survivorship curves and imaging of five of six different biological replicates of LL vs SL worms that was used for RNA-Seq analysis (Replicate 2 shown in Figure 1). B-E. Validation of gene expression changes in LL vs SL worms in RNA-Seq. qRT-PCR of metabolic genes B. *fat-7* C. *acdh-1* D. *acdh-2* and E. *gst-10* in the SL and LL worm sub-populations at Day 6 (\*\*\*\**P*≤0.0001, \*\*\**P*≤0.001, \*\**P*≤0.01, \**P*≤0.05; ns P>0.05). *P*-values calculated with unpaired, two-tailed Welch’s t-test. qRT–PCR data are mean + s.e.m. of 3 biological replicates. F. WormCat visualization of categories enriched in genes that exhibit differential local splicing events and differential transcript usage in LL vs SL worms.

**Figure S2:**
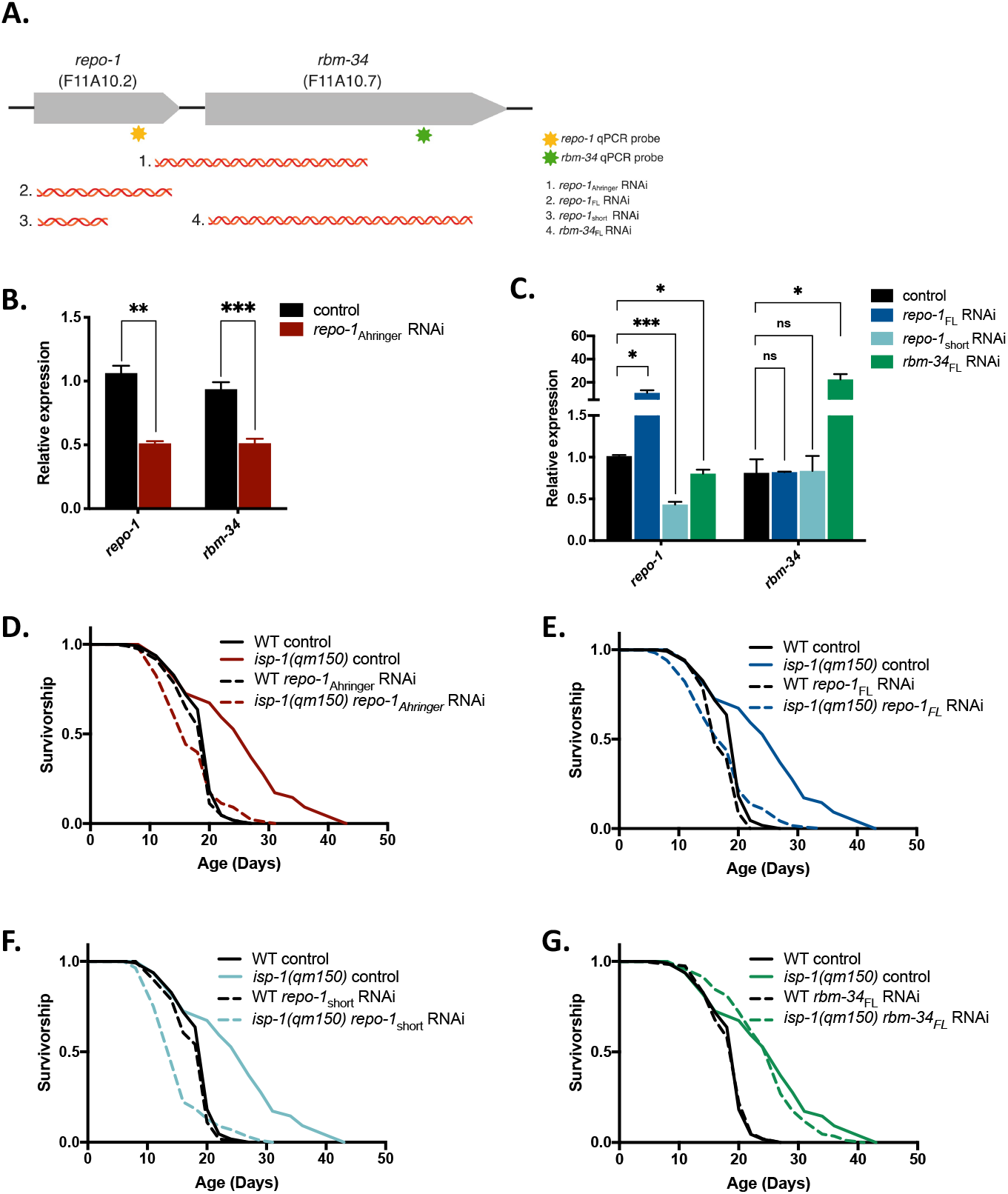
Specificity of RNAi to REPO-1. A. Schematic representing the position of genes *repo-1* and *rbm-34* in the *C. elegans* genome operon CEOP4488 and the target sequences of the old *repo-1* RNAi from the Ahringer library (*repo-1*_Ahringer_ RNAi) and the newly constructed *repo-1*_FL_, *repo-1*_short_ RNAi and *rbm-34*_FL_ RNAi. Approximate positions of the qRT-PCR probes are marked. B. *repo-1*_Ahringer_ RNAi knocks down both *repo-1* and *rbm-34*. qRT-PCR of *repo-1* and *rbm-34* expression in Day 1 wildtype worms on control and *repo-1*_Ahringer_ RNAi from hatch. C. qRT-PCR of *repo-1* and *rbm-34* expression in Day 1 wildtype worms fed with control, *repo-1*_FL_, *repo-1*_short_ and *rbm-34*_FL_ RNAi from hatch (\*\*\*\**P*≤0.0001, \*\*\**P*≤0.001, \*\**P*≤0.01, \**P*≤0.05; ns P>0.05). *P*-values calculated with unpaired, two-tailed Welch’s t-test. qRT–PCR data are mean + s.e.m. of 3 biological replicates. Increased signal of *repo-1* and *rbm-34* on treatment with *repo-1*_FL_ and *rbm-34*_FL_ RNAi respectively is likely due to non-specific signals from the siRNAs produced by bacteria that are ingested by the worms. Note that *repo-1*_short_ RNAi is able to knockdown *repo-1* with equal efficiency as *repo-1*_Ahringer_ RNAi but has no effect on *rbm-34* expression. D-G. Knockdown of *repo-1* and not *rbm-34* blocks lifespan extension in a C. elegans model of longevity. Survivorship of wildtype (WT) and *isp-1(qm150)* worms with or without D. *repo-1*_Ahringer_ (*P*=0.5434), E. *repo-1*_FL_ (*P*=0.1228), F. *repo-1*_short_ RNAi (*P*=0.0006, shorter-lived) and G. *rbm-34*_FL_ RNAi (*P*= <0.0001) (*P*-values are comparing wildtype RNAi vs *isp-1(qm150)* RNAi in each case).

**Figure S3:**
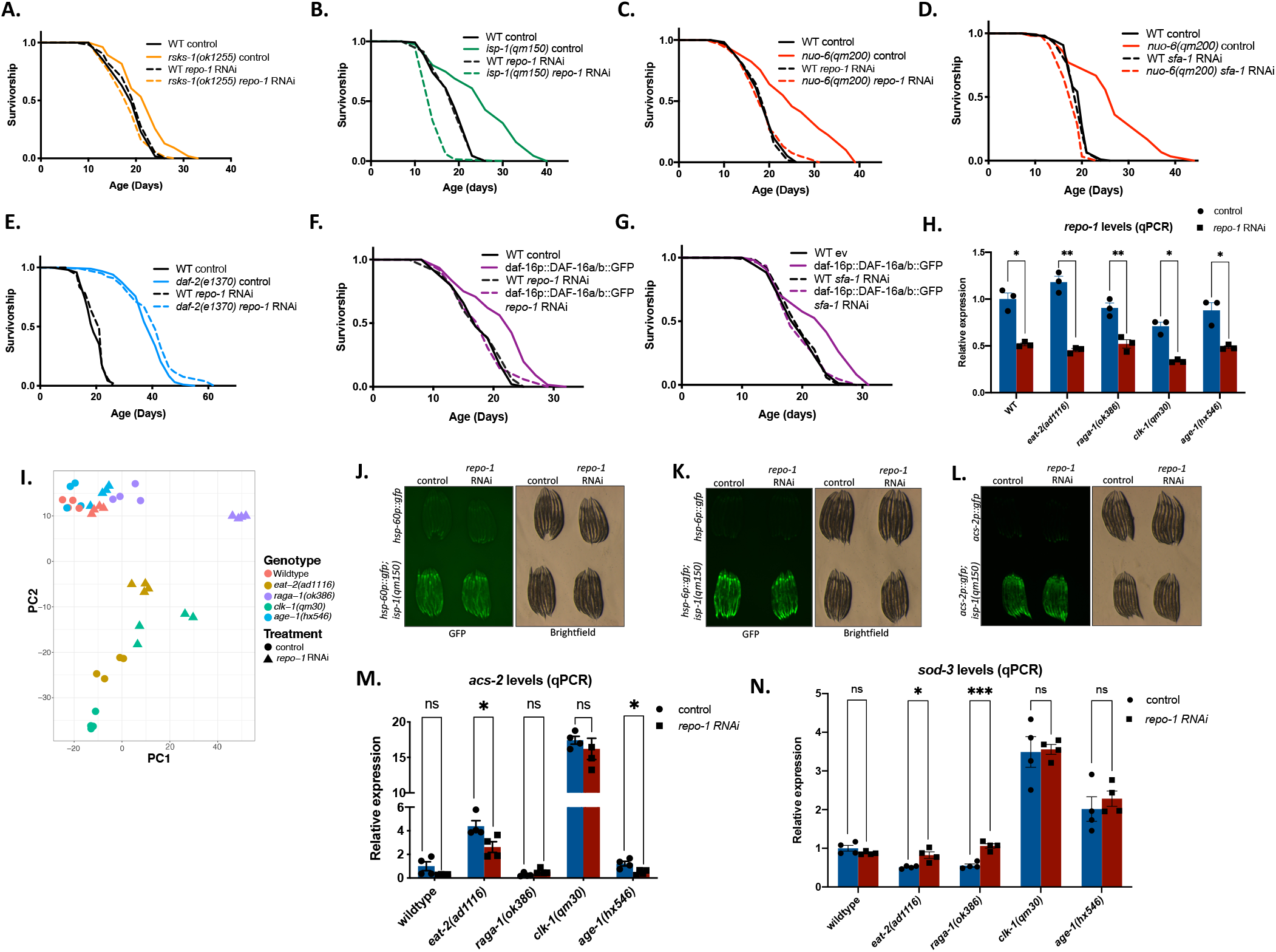
The effect of loss of REPO-1 and SFA-1 is conserved across other mutants in the different longevity pathways. A. Effect of loss of REPO-1 is conserved across other mutants of the TORC1 pathway. Survivorship of wild-type (WT) and *rsks-1(ok1255)* -/+ *repo-1* RNAi *(P=0*.*0803*, wildtype N2 *repo-1* RNAi versus *rsks-1(ok1255) repo-1* RNAi, 2 replicates). B-D. Effect of loss of REPO-1 is conserved across other mutants of the ETC pathway. B. Survivorship of wild-type (WT) and *isp-1(qm50)* on *repo-1* RNAi *(P<0*.*0001*, wildtype N2 *repo-1* RNAi versus *isp-1(qm50) repo-1* RNAi, 2 replicates). C. Survivorship of WT and *nuo-6(qm200)* on *repo-1* RNAi *(P=0*.*9116*, wildtype N2 *repo-1* RNAi versus *nuo-6(qm200) repo-1* RNAi, 2 replicates). D. Survivorship of WT and *nuo-6(qm200)* on *sfa-1* RNAi *(P<0*.*0001*, wildtype N2 *sfa-1* RNAi versus *nuo-6(qm200) sfa-1* RNAi, 2 replicates). E. Effect of loss of REPO-1 is conserved across other mutants of the rIIS pathway. Survivorship of wild-type (WT) and *daf-2(e1370) -/+ repo-1* RNAi *(P<0*.*0001*, wildtype N2 *repo-1* RNAi versus *daf-2(e1370) repo-1* RNAi, 2 replicates). F, G. Survivorship of wild-type (WT) and DAF-16a/b overexpressor worms (*daf-16p*::DAF16a/b::GFP) *-/+ repo-1* RNAi (*P*=0.8994) and *sfa-1* RNAi (*P*=0.8718) (*P*-values are comparing WT on RNAi vs mutant on RNAi, 2 replicates). H. REPO-1 is knocked down in different mutants with equal efficiency. qPCR showing ∼50% knockdown of *repo-1* in different longevity mutants fed with empty vector and *repo-1* RNAi from hatch and collected at Day 1 of adulthood (\*\*\*\**P*≤0.0001, \*\*\**P*≤0.001, \*\**P*≤0.01, \**P*≤0.05; ns P>0.05). *P*-values calculated with unpaired, two-tailed Welch’s t-test. qRT–PCR data are mean + s.e.m. of 3 biological replicates. I. Principal Component Analysis of RNA Seq Samples in different longevity mutants -/+ *repo-1* RNAi. J,K. Knockdown of REPO-1 does not abrogate mitoUPR in ETC mutants. Transcriptional induction of mitochondrial chaperones *hsp-6p::GFP and hsp-60p::GFP* in wildtype N2 and ETC mutant *isp-1(qm150)* without or with *repo-1* RNAi from hatch and imaged at Day 1 of adulthood. L. Knockdown of REPO-1 does not abrogate *acs-2* induction in ETC mutants. Day 1 imaging of *acs-2p::GFP* in wildtype N2 and *isp-1(qm150)* background without or with *repo-1* RNAi from hatch. M, N. RT-qPCR of *acs-2* and *sod-3* in Day 1 WT and longevity mutants fed on control bacteria or *repo-1* RNAi from hatch. (\*\*\*\**P*≤0.0001, \*\*\**P*≤0.001, \*\**P*≤0.01, \**P*≤0.05; ns P>0.05). *P*-values calculated with unpaired, two-tailed Welch’s t-test. qRT–PCR data are mean + s.e.m. of 4 biological replicates. Samples here are the same as used in RNA-Seq (Fig 2).

**Fig S4:**
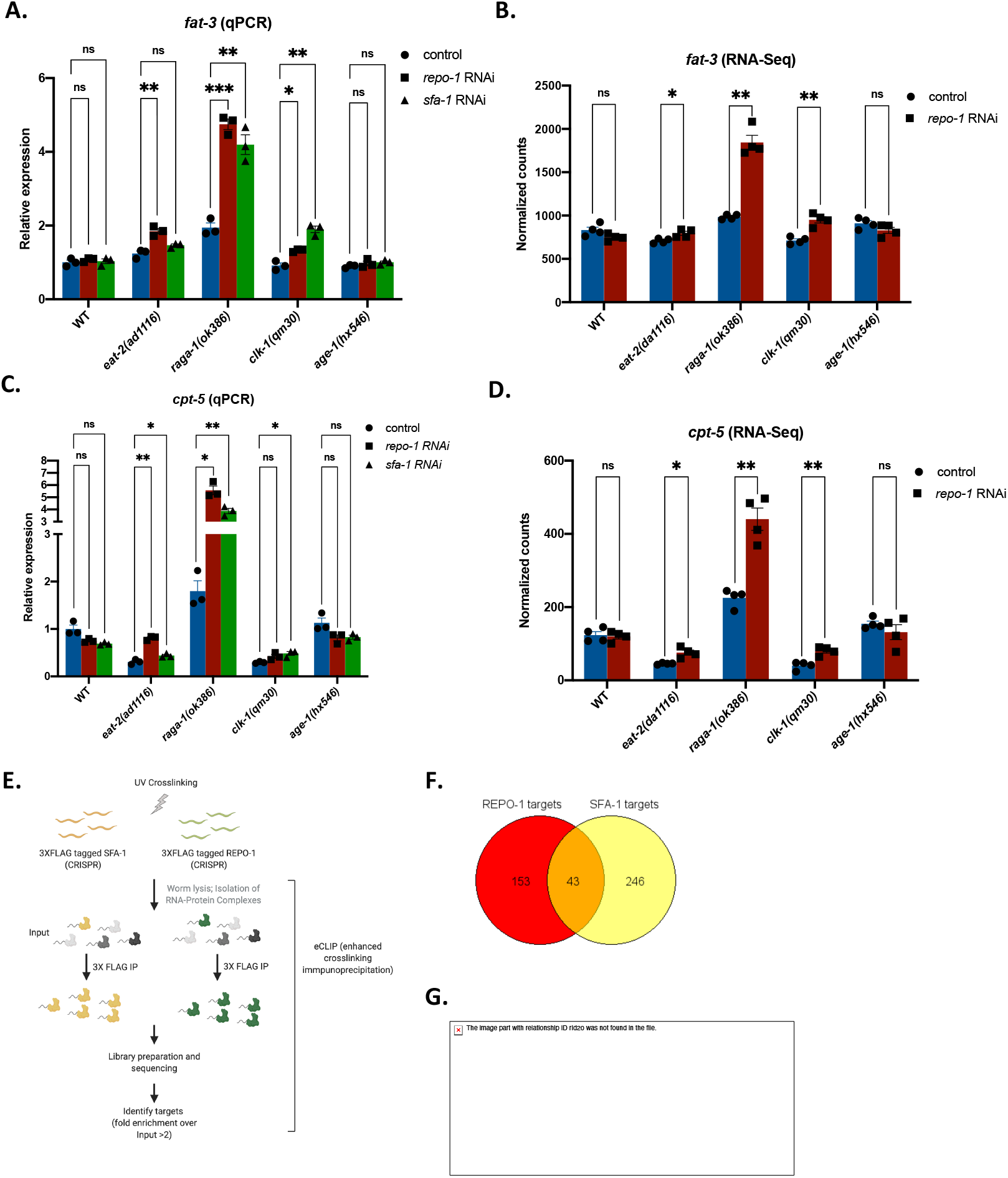
Loss of REPO-1 specifically affects lipid metabolism in splicing factor sensitive longevity pathways. A. Expression of *fat-3* by qRT-PCR in wildtype and different mutants - /+ *repo-1* and *sfa-1* RNAi at Day 1. B. Normalized counts of *fat-3* transcript in RNA-Seq of wildtype and different mutants -/+ *repo-1* RNAi at Day 1. C. Expression of *cpt-5* by qRT-PCR in wildtype and different mutants -/+ *repo-1* and *sfa-1* RNAi at Day 1. D. Normalized counts of *cpt-5* transcript in RNA-Seq of wildtype and different mutants -/+ *repo-1* RNAi at Day 1. (\*\*\*\**P*≤0.0001, \*\*\**P*≤0.001, \*\**P*≤0.01, \**P*≤0.05; ns P>0.05). *P*-values calculated with unpaired, two-tailed Welch’s t-test. RNA-seq data are mean + s.e.m. of normalized read counts of 4 biological replicates. qRT–PCR data are mean + s.e.m. of 3 biological replicates. E. Schematic representing eCLIP in worms. F. Venn diagram displaying the overlap of RNA targets with enrichment >2 fold (log2 fold change >1, IP vs Input) in SFA-1 and REPO-1. G. Validation of *tos-1* as a target of SFA-1 and REPO-1. Semi-quantitative PCR showing differential isoforms of *tos-1* in Day 1 worms on loss of *repo-1* and *sfa-1* using RNAi. Red arrow marks the appearance of a different isoform of *tos-1* on loss of *repo-1* and *sfa-1*.

**Figure S5:**
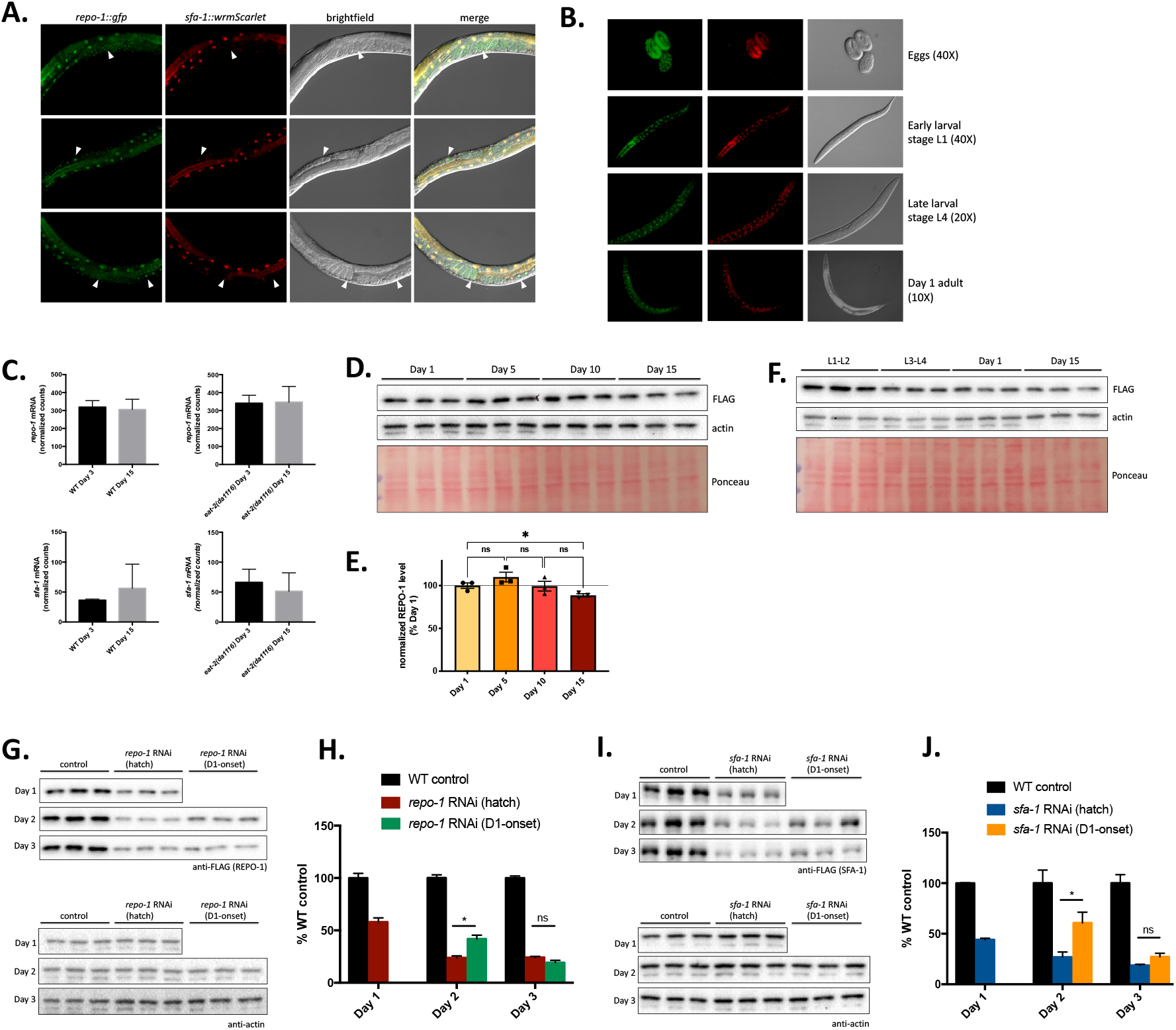
REPO-1 is expressed at elevated levels in early larval stages of worm development and might be crucial for its function in this time window. A. Fluorescent microscope images of worms with CRISPR tagged REPO-1::GFP and CRISPR tagged SFA-1::wrmScarlet at Day 1. Worms imaged at 20X magnification. White arrows mark the presence of REPO-1 and absence of SFA-1 in early embryos. B. Fluorescent microscope images of CRISPR tagged REPO-1 fused to GFP and CRISPR tagged SFA-1 fused to wrmScarlet from the egg stage to Day 1 of adulthood. C. REPO-1 and SFA-1 mRNA levels do not change with age. Normalized counts of *repo-1* and *sfa-1* transcripts at Day 3 and Day 15 in WT and *eat-2(ad1116)* worms. Transcript counts obtained from previously published RNA-Seq data (Heintz et al. Nature 2017). D,E. REPO-1 protein levels do not change with age. D. Western blotting of CRISPR tagged endogenous 3XFLAG::REPO-1 worms at Day1, Day 5, Day 10 and Day 15 of adulthood. Blots probed with 3XFLAG and actin antibodies. Ponceau staining of the blot shows equal loading. E. Quantification of 3XFLAG:REPO-1 normalized to actin. Blots quantified using ImageJ and represented as percent of expression at Day 1 of adulthood. F. Western blotting of CRISPR tagged endogenous 3XFLAG::REPO-1 worms at L1-L2, L3-L4, Day 1 and Day 15 of adulthood. Blots probed with 3XFLAG and actin antibodies. Ponceau staining of the blot shows equal loading. G-J. Adult-onset RNAi efficiently knocks down REPO-1 and SFA-1 comparable to RNAi from hatch. Western blotting of CRISPR tagged endogenous G. 3XFLAG::REPO-1 worms -/+ *repo-1* RNAi; I. 3XFLAG::SFA-1 worms -/+ *sfa-1* RNAi from hatch or Day 1 of adulthood (D1-onset). Samples collected on Day 1, Day 2 and Day 3 to measure efficiency of knockdown. Lysates probed for 3XFLAG as a readout of REPO-1/SFA-1 and actin as loading control. Quantification of knockdown of H. REPO-1 and J. SFA-1 normalized to actin. Blots quantified using ImageJ and represented as percent of RNAi untreated control. (\*\*\*\**P*≤0.0001, \*\*\**P*≤0.001, \*\**P*≤0.01, \**P*≤0.05; ns P>0.05). *P*-values calculated with unpaired, two-tailed Welch’s t-test.

## Notes

### Competing Interest Statement

The authors have declared no competing interest.

